# Protein phosphatases regulate the formation of Müller glia-derived progenitor cells in the chick retina

**DOI:** 10.1101/2023.12.11.570629

**Authors:** Lisa E. Kelly, Heithem M. El-Hodiri, Andrew Crider, Andy J. Fischer

## Abstract

Different kinase-dependent cell signaling pathways are known to play important roles in glia-mediated neuroprotection and reprogramming of Müller glia (MG) into Müller glia-derived progenitor cells (MGPCs) in the retina. However, very little is known about the phosphatases that regulate kinase-dependent signaling in MG. Using single-cell RNA-sequencing (scRNA-seq) databases, we investigated patterns of expression of Dual Specificity Phosphatases (DUSP1/6) and other protein phosphatases in normal and damaged chick retinas. We found that *DUSP1, DUSP6, PPP3CB, PPP3R1* and *PPPM1A/B/D/E/G* are dynamically expressed by MG and MGPCs in retinas during the process of reprogramming. We find that inhibition of DUSP1/6 and PP2C phosphatases enhances the formation of proliferating MGPCs in damaged retinas and in retinas treated with insulin in FGF2 in the absence of damage. By contrast, inhibition of PP2B phosphatases suppressed the formation of proliferating MGPCs, but increased numbers of proliferating MGPCs in undamaged retinas treated with insulin and FGF2. In damaged retinas, inhibition of DUSP1/6 increased levels of pERK1/2 and cFos in MG whereas inhibition of PP2B’s decreased levels of pStat3 and pS6 in MG. Analyses of scRNA-seq libraries identified numerous differentially activated gene modules in MG in damaged retinas versus MG in retinas treated with insulin+FGF2 suggesting significant differences in kinase-dependent signaling pathways that converge on the formation of MGPCs. Inhibition of phosphatases had no significant effects upon numbers of dying cells in damaged retinas. We conclude that the activity of different protein phosphatases “fine-tune” the cell signaling responses of MG in damaged retinas and during the reprogramming of MG into MGPCs.

## Introduction

Müller glia (MG) are the predominant type of support cell in the retina that are common to all vertebrate classes. MG perform many important glial functions including structural support, synaptic support, osmotic homeostasis, and metabolic support. However, MG also have the extraordinary capability to de-differentiate, proliferate, acquire progenitor phenotype, and generate new neurons [1–5]. Reprogramming of MG involves activation, down-regulation of glial genes, up-regulation of progenitor-associated factors, proliferation, followed by neuronal differentiation (Hoang et al., 2020). The capacity of MG to regenerate retinal neurons varies greatly between vertebrate classes. The MG in fish retinas have an extraordinary ability to effectively become proliferating progenitor cells that produce functional neurons to restore vision to damaged retinas (reviewed by [6, 7]). The MG in mouse retinas respond to damage by becoming reactive and activating networks of genes to restore quiescence [8]. The regenerative capacity of MG in chick retinas lies somewhere in between that of fish and mammals. MG in the chick retina form numerous proliferating MGPCs with limited neurogenic potential in response to acute neuronal damage or in the absence of damage when treated with insulin and FGF2 [2, 9].

In the mouse model, recent studies indicated that forced-expression of the transcription factor *Ascl1*, in combination with NMDA-induced damage and HDAC-inhibitor, stimulates reprogramming of MG into functional, light-responsive neurons [10–12]. The numbers of neurons produced by Ascl1-over expressing MG can be enhanced by inhibiting Jak/Stat [13], ablating microglia from the retina [14], inhibiting NFkB, inhibiting ID transcription factors, inhibiting TGFβ/Smad3 [15] and enhanced by combined forced expression of Ascl1 and Atoh1 [16]. Cell signaling cascades are broadly and rapidly activated in MG during the process of reprogramming. Pathways that promote the formation of MGPCs include MAPK [17, 18], Jak/Stat [19, 20], Wnt/β-catenin [21–23], Hedgehog [24, 25], Notch [26, 27], retinoic acid [28], BMP/Smad1/5/8 [29], cannabinoid [30], midkine [31] and mTor [32]. Pathways that suppress the formation of MGPCs include glucocorticoid [33], TGFβ/Smad2/3 [29, 34] and NFkB [15, 35]. The majority of these cell signaling pathways involve kinase-dependent cascades, and it is assumed these pathways are negatively regulated by phosphatases. The purpose of this study was to understand how phosphatases regulate the responses of MG to damage and become proliferating progenitor-like cells. Accordingly, we investigated patterns of expression of different protein phosphatases in MG following neuronal damage or treatment with insulin and FGF2, and how inhibition of different families of phosphatases influence glial de-differentiation and the proliferation of MGPCs *in vivo*.

## Methods and Materials

### Animals

The animals approved for use in these experiments were in accordance with the guidelines established by the National Institutes of Health and IACUC at The Ohio State University. Newly hatched P0 wildtype leghorn chicks (*Gallus gallus domesticus*) were obtained from Meyer Hatchery (Polk, Ohio). Post-hatch chicks were maintained in a regular diurnal cycle of 12 hours light, 12 hours dark (8:00 AM-8:00 PM). Chicks were housed in stainless-steel brooders at 25°C and received water and Purina^tm^ chick starter *ad libitum*.

### Intraocular injections

Chicks were anesthetized with 2.5% isoflurane mixed with oxygen from a non-rebreathing vaporizer. The technical procedures for intraocular injections were performed as previously described [36]. With all injection paradigms, pharmacological and vehicle treatments were administered to the right and left eye respectively. Compounds were injected in 20 μl sterile saline with 0.05 mg/ml bovine serum albumin added as a carrier. Compounds included: NMDA (38.5nmol or 154 µg/dose; Sigma-Aldrich), FGF2 (250 ng/dose; R&D systems). 5-Ethynyl-2-deoxyuridine (EdU) was intravitreally injected to label the nuclei of proliferating cells. Injection paradigms are included in each figure.

### Preparation of clodronate liposomes

Clodronate liposomes were synthesized utilizing a modified protocol from previous descriptions [37–39]. In short, 8 mg of L-α-Phosphatidyl-DL-glycerol sodium salt (Sigma P8318) was dissolved in chloroform. 50 mg of cholesterol was dissolved in chloroform with the lipids in a microcentrifuge tube. This tube was rotated under nitrogen gas to evaporate the chloroform and leave a thin lipid-film on the walls of the tube. 158 mg dichloro-methylene diphosphonate (clodronate; Sigma-Aldrich) dissolved sterile PBS (pH 7.4) was added to the lipid/cholesterol film and vortexed for 5 minutes. To reduce size variability of lipid vesicles, the mixture was sonicated at 42 kHz for 6 minutes. Liposomes were purified via centrifugation at 10,000 x G for 15 minutes, supernatant was aspirated, and liposomes resuspended in 150 µl PBS. Each intravitreal injection of clodronate-liposome solution was between 5 and 20 µl. The variable yield of clodronate-liposomes during the purification resulted in variable efficacy per dose. Accordingly, the dosage was adjusted such that >98% of the microglia are ablated by 2 days after administration with no off-target cell death.

### Single Cell RNA sequencing

We analyzed scRNA-seq libraries that were generated and described previously [8, 30, 30]. To improve gene annotation and improve cell calling, we re-derived scRNA-seq libraries using updated genome reference libraries and updated 10X Cell Ranger software. In short, FASTQ files were regenerated from BAM files using bam2fastq (CellRanger 7.0.1). BAM files used were from two replicates each of scRNA-seq librarties prepared from saline- or NMDA-treated chick retina or from scRNA-seq libraries from chick retinas treated with 2 or 3 daily doses of FGF and insulin. Regenerated FASTQ files were processed as described previously using CellRanger count and chicken reference package prepared using CellRanger mkref (CellRanger 7.0.1). Chicken reference package was prepared from FASTA and GTF files ((Ensembl.org, bGalGal1.mat.broiler.GRC7b v 108).

Gene expression was counted using unique molecular identifier bar codes, and gene-cell matrices were constructed. Using Seurat toolkits, Uniform Manifold Approximation and Projection for Dimension Reduction (UMAP) plots were generated from aggregates of multiple scRNA-seq libraries [40, 41]. Seurat was used to generate lists of differentially expressed genes (DEGs), heatmap feature plots and dot plots. Significance of difference in was determined using a Wilcoxon Rank Sum test with Bonferroni correction. Genes that were used to identify different types of retinal cells included the following: (1) Müller glia: *GLUL, VIM, SCL1A3, RLBP1*, (2) MGPCs: *PCNA, CDK1, TOP2A, ASCL1*, (3) microglia: *C1QA, C1QB, CCL4, CSF1R, TMEM22*, (4) ganglion cells: *THY1, POU4F2, RBPMS2, NEFL, NEFM*, (5) amacrine cells: *GAD67, CALB2, TFAP2A*, (6) horizontal cells: *PROX1, CALB2, NTRK1*, (7) bipolar cells: *VSX1, OTX2, GRIK1, GABRA1*, and (7) cone photoreceptors: *CALB1, GNAT2, OPN1LW*, and (8) rod photoreceptors: *RHO, NR2E3, ARR3.* The MG have an over-abundant representation in the scRNA-seq databases. This likely resulted from fortuitous capture-bias and/or tolerance of the MG to the dissociation process.

Gene Ontology (GO) enrichment analysis was performed using ShinyGO V0.72 (http://bioinformatics.sdstate.edu/go/). Adjustment for multiple hypothesis testing is performed through ShinyGo by normalizing the enrichment score for each gene set to account for the size of the set, to produce a normalized enrichment score (NES). The proportion of false positives is controlled by calculating the false discovery rate (FDR) corresponding to each NES. The FDR enrichment p-value is calculated as the probability that a gene set with a given NES represents a false positive finding; the probability is computed by comparing the tails of the observed and null distributions for the NES [42].

CellRanger output files for Gene-Cell matrices for scRNA-seq data for libraries from saline and NMDA-treated retinas are available through GitHub: https://github.com/jiewwwang/Singlecell-retinal-regeneration or Sharepoint: chick embryonic retina scRNA-seq Cell Ranger outs (https://osumc.sharepoint.com/:f:/s/Links/Eoto-Qg2uuxDn1bHWMM6gdkBTft4S_YSBjQJResxY-qehA?e=mdaPlg) and [chick retina scRNA-seq Cell Ranger output files](https://osumc.sharepoint.com/sites/Links/FischerLab/Forms/AllItems.aspx?id=%2Fsites%2FLinks%2FFischerLab%2Fchick%20embryonic%20retina%20scRNA%2Dseq%20Cell%20Ranger%20outs&p=true&ga=1) (https://osumc.sharepoint.com/:f:/s/Links/Eoto-Qg2uuxDn1bHWMM6gdkBTft4S_YSBjQJResxY-qehA?e=mdaPlg). scRNA-seq datasets are deposited in GEO (GSE135406, GSE242796) and Gene-Cell matrices for scRNA-seq data for libraries from saline and NMDA-treated retinas are available through NCBI (GSM7770646, GSM7770647, GSM7770648, GSM7770649).

### Fixation, sectioning and immunocytochemistry

Retinal tissue samples were formaldehyde fixed, sectioned, and labeled via immunohistochemistry as described previously[43, 44]. Antibody dilutions and commercial sources for images used in this study are described in table 1. Observed labeling was not due to off-target labeling of secondary antibodies or tissue autofluorescence because sections incubated exclusively with secondary antibodies were devoid of fluorescence. Secondary antibodies utilized include donkey-anti-goat-Alexa488/568, goat-anti-rabbit-Alexa488/568/647, goat-anti-mouse-Alexa488/568/647, goat-anti-rat-Alexa488 (Life Technologies) diluted to 1:1000 in PBS and 0.2% Triton X-100.

**Table 1.** List of antibodies, working dilution, clone/catalog number and source.

| Antibody | Dilution | Host | Clone/Catalog number | Source |
| --- | --- | --- | --- | --- |
| Sox2 | 1:1000 | Goat | KOY0418121 | R&D |
| Sox9 | 1:2000 | Rabbit | AB5535 | Millipore |
| pERK1/2 | 1:600 | Rabbit | 4370S | Cell Signaling |
| pS6 | 1:750 | Rabbit | #2215; Ser240/244 | Cell Signaling |
| pStat3 | 1:300 | Rabbit | 9131 | Cell Signaling |
| pSmad1/5/8 | 1:200 | Rabbit | D5B10 | Cell Signaling |
| cFos | 1:200 | Rabbit | SC-253 | Santa Cruz |
| Anti-goat IgG Alex488 | 1:1000 | Donkey | A11055 | Invitrogen |
| Anti-goat IgG Alex568 | 1:1000 | Donkey | A11057 | Invitrogen |
| Anti-rabbit IgG Alex488 | 1:1000 | Donkey | A21206 | Invitrogen |
| Anti-rabbit IgG Alex568 | 1:1000 | Donkey | A10042 | Invitrogen |
| Anti-rabbit IgG Alex647 | 1:1000 | Goat | A32733 | Invitrogen |
| Anti-mouse IgG Alexa Fluor 488 | 1:1000 | Donkey | A21202 | Invitrogen |
| Anti-mouse IgG Alexa Fluor 594 | 1:1000 | Donkey | A32744 | Invitrogen |

### Fluorescent in situ hybridization (FISH)

The FISH protocol has been adapted from the hybridization chain reaction (HCR) for tissue sections provided by Molecular Instruments. In short, tissues were dissected and fixed in 4% PFA in DEPC water with 2mM EDTA for 4 hrs. Tissues were washed for 30 min in 0.1% Tween-20 PBS (PWT) and soaked overnight into 30% sucrose. Tissues were cryosectioned and thaw-mounted onto Fisherbrand^tm^ Superfrost^tm^ Plus slides (Fisher Scientific). Retinal sections were rehydrated in PWT for 15 min followed by 2x SSCT washes for 20 minutes. Slides were added with hybridization buffer and incubated at 37°C. The buffer was aspirated and the probe-set diluted 1:50 in hybridization buffer and incubated for 24hrs in a humidified chamber at 37°C. Probes were aspirate, tissues were incubated in hybridization buffer for 15 minutes at 37°C, and washed in 5x SSCT for 5 minutes at room temperature. Amplification primers were snap cooled, mixed, and applied to the sections at room temperature for 24hrs in the dark. The primers were washed off with 5x SSCT for 10 minutes, and glass cover slips mounted for imaging.

### Labeling for EdU

For the detection of nuclei that incorporated EdU, immunolabeled sections were fixed in 4% formaldehyde in 0.1M PBS pH 7.4 for 5 minutes at room temperature. Samples were washed for 5 minutes with PBS, permeabilized with 0.5% Triton X-100 in PBS for 1 minute at room temperature and washed twice for 5 minutes in PBS. Sections were incubated for 30 minutes at room temperature in a buffer consisting of 100 mM Tris, 8 mM CuSO_4_, and 100 mM ascorbic acid in dH_2_O. The Alexa Fluor 568 Azide (Thermo Fisher Scientific) was added to the buffer at a 1:100 dilution.

### Terminal deoxynucleotidyl transferase dUTP nick end labeling (TUNEL)

The TUNEL assay was implemented to identify dying cells by imaging fluorescent labeling of double stranded DNA breaks in nuclei. The *In Situ* Cell Death Kit (TMR red; Roche Applied Science) was applied to fixed retinal sections as per the manufacturer’s instructions.

### Photography, measurements, cell counts and statistics

Microscopy images of retinal sections were captured with the Leica DM5000B microscope with epifluorescence and the Leica DC500 digital camera. High resolution confocal images were obtained with a Leica SP8 available in The Department of Neuroscience Imaging Facility at The Ohio State University. Representative images are modified to have enhanced color, brightness, and contrast for improved clarity using Adobe Photoshop. For quantification of numbers of EdU^+^ cells, a fixed region of retina was counted and average numbers of Sox2^+^/CD45^+^ and EdU labeled cells. The retinal region selected cell counts was standardized between treatment and control groups to reduce variability and improve reproducibility.

Similar to previous reports [29, 45, 46], immunofluorescence was quantified by using Image J (NIH). Identical illumination, microscope, and camera settings were used to obtain images for quantification. Retinal areas were sampled from 5.4 MP digital images. These areas were randomly sampled over the inner nuclear layer (INL) where the nuclei of the bipolar and amacrine neurons were observed. The density sum was calculated as the total of pixel values for all pixels within regions above threshold. The mean density sum was calculated for the pixels within threshold regions from ≥5 retinas for each experimental condition. GraphPad Prism 6 was used for statistical analyses.

Measurement for immunofluorescence of pERK1/2, pStat3 and pS6 in MG/MGPCs using Image J, as described previously [29, 45–47]. In short, regions of interest were identified in single optical confocal sections by selecting the total area of pixel values above threshold (≥70) for Sox2 or Sox9 immunofluorescence (in the red channel) and copying nuclear cFos or pStat3 from only MG (in the green channel). Measurements of pS6 or pERK1/2 immunofluorescence were made for a fixed, cropped area (14,000 µm^2^) of INL, OPL and ONL. Measurements were made for regions containing pixels with intensity values of 70 or greater (0 = black and 255 = saturated). The intensity sum was calculated as the total of pixel values for all pixels within threshold regions. The mean intensity sum was calculated for the pixels within threshold regions from ≥6 retinas for each experimental condition.

For statistical evaluation of differences in treatments, a two-tailed paired *t*-test was applied for intra-individual variability where each biological sample also served as its own control. For two treatment groups comparing inter-individual variability, a two-tailed unpaired *t*-test was applied. For multivariate analysis, an ANOVA with the associated Tukey Test was used to evaluate any significant differences between multiple groups.

## Results

### Expression of protein phosphatases in mature and damaged retinas

We began by probing for patterns of expression of different protein phosphatases in a scRNA-seq database that was generated for early time-points, 3, 12 and 48 hours, after NMDA-treatment, as originally described in detail in previous studies [30, 48]. UMAP plots were generated and clusters of different cells were identified based on well-established patterns of expression (Fig. 1a,b). For example, resting MG formed a discrete cluster of cells and expressed *GLUL* and *RLBP1* (Fig. 1c). After damage, MG down-regulate markers of mature glia as they transition into reactive glial cells and up-regulate genes such as *TGFB2* and *PMP2* (Fig. 1d). MG forming distinct clusters based on the time after NMDA-treatment (Fig. 1a-d). *DUSP1* was widely detected in all types of retinal neurons and glia, with scattered expression in RGCs (Fig. 1e). By comparison, *DUSP6* was not detected in oligodendrocytes, NIRG cells, or RGCs, but was detected in cone photoreceptors, rod photoreceptors, HCs, and scattered BPs, ACs and MG (Fig. 1f). *PPP3CB, PPP3R1, PPM1A, PPM1B, PPM1C, PPM1D, PPM1E* and *PPM1G* were widely expressed by all types of retinal neurons and glia, with the exception of oligodendrocytes and NIRG cells (Fig. 1g-m). We bioinformatically isolated MG and performed a more detailed analyses of expression patterns of protein phosphatases based on treatment/UMAP ordering. All protein phosphatases were rapidly and significantly (p<0.0001) upregulated in MG by 3hrs after NMDA, with the exception of *DUSP1* which was upregulated by 12hrs after NMDA (Fig. 1n; Supplemental table 1). All protein phosphatases were significantly downregulated by MG at 48hrs after NMDA compared to earlier times after NMDA (Fig. 1n; Supplemental table 1).

**Figure 1.**
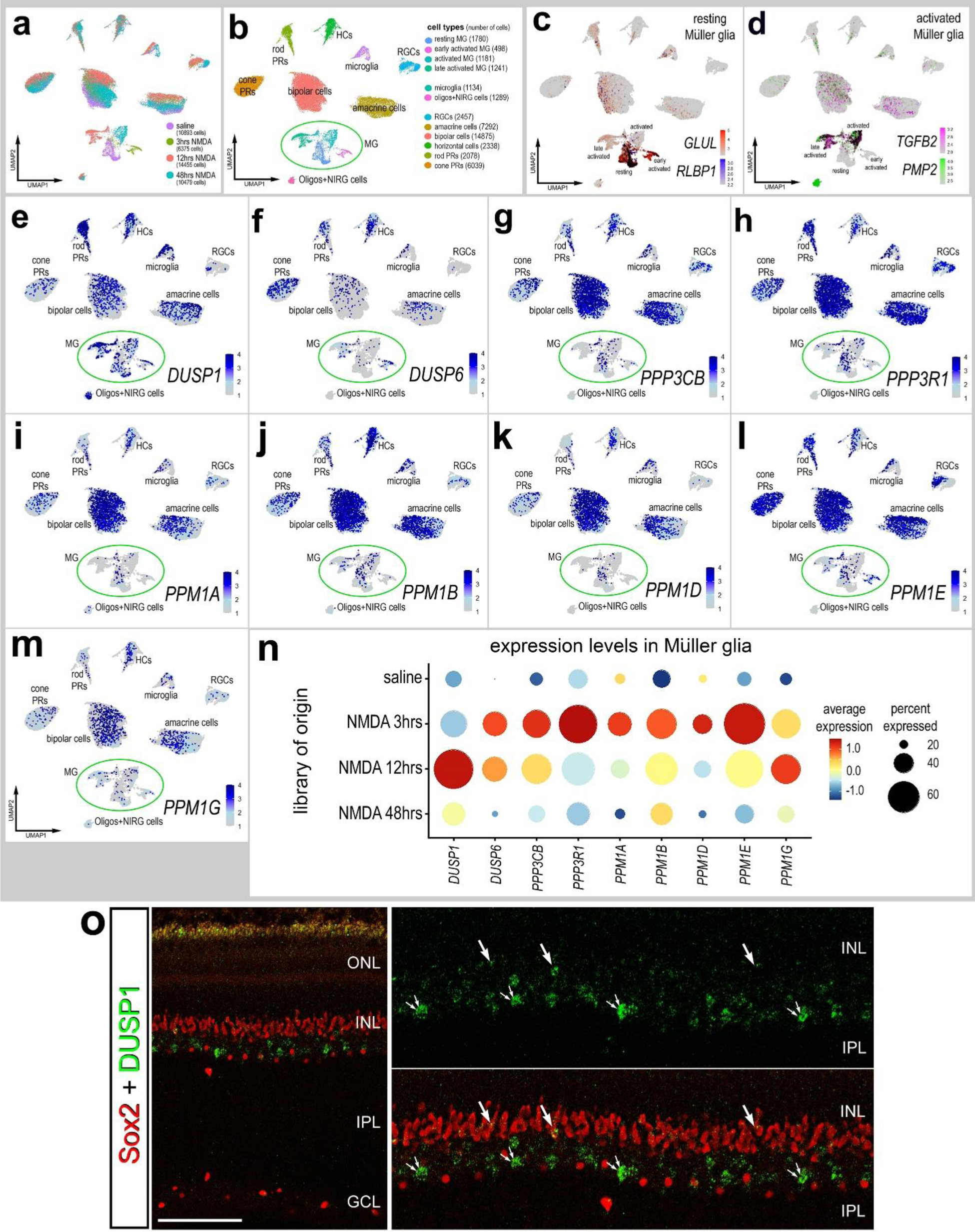
Expression of Phosphatase-related genes in chick retinal cells. scRNA-seq was used to identify patterns and levels of expression of protein phosphatases in control and NMDA-damaged retinas at 3, 12 and 48 hours after treatment (**a**). UMAP clusters of cells were identified based on well-established patterns of gene expression (see Methods; **b**). MG were identified by expression of *RLBP1* and *GLUL* in resting MG (**c**), and *TGFB2* and *PMP2* in activated MG (**d**). Each dot represents one cell and black dots indicate cells that express 2 or more genes (**c,d**). UMAP heatmap plots (**e-m**) illustrate patterns and levels of expression for phosphatases in retinal neurons and glia. The dot plots in **n** illustrates average expression (heatmap) and percent expressed (dot size) for different phosphatases in MG that are significantly (p<0.0001) up- or downregulated at different times following NMDA-treatment. Significance of difference was determined by using a Wilcox rank sum test with Bonferroni correction. Fluorescence *in situ* hybridization (**o**): Retinal sections were labeled with antibodies to Sox2 and fluorescence *in situ* hybridization for *DUSP1*. Arrows indicate MG and small double arrows indicate amacrine cells. Abbreviations: ONL – outer nuclear layer, INL – inner nuclear layer, IPL – inner plexiform layer, GCL – ganglion cell layer. The calibration bar represents 50 µm (**o**).

Patterns of expression of *DUSP1* closely matched between scRNA-seq libraries and fluorescence *in situ* hybridization (FISH). FISH signal for *DUSP1* was observed in some MG and presumptive amacrine cells in the inner INL (Fig. 1o), similar to patterns of expression seen in scRNA-seq libraries (Fig. 1e). However, unlike patterns of expression seen in scRNA-seq libraries, very little FISH signal was detected in rod photoreceptors in the ONL or distal INL in putative bipolar and horizontal cells (Fig. 1o).

### Inhibition of protein phosphatases in damaged retinas

We applied different small molecule inhibitors to target different families of protein phosphatases. FK506 is known to inhibit the PP2B family of phosphatases which includes PPP3CB and PPP3R1 regulatory subunit of calcineurin [49, 50]. BCI is known to inhibit DUSP1 and DUSP6 [51]. Sanguinarine is known to inhibit PP2C family of Mg2^+^/Mn2^+^ -dependent phosphatases which includes PPM1A, PPM1B, PPM1C, PPM1D, PPM1E and PPM1G [52–54]. We found that injection of FK506 significantly reduced numbers of proliferating Sox2^+^ MGPCs om damaged retinas (Fig. 2a,b). BCI had no significant effect upon the formation of proliferating MGPCs in central regions of the retina, but significantly increased numbers of proliferating MGPCs in peripheral regions of damaged retinas (Fig. 2c,d). Similar to the effects of BCI, treatment with sanguinarine significantly increased numbers of proliferating Sox2^+^ MGPCs in damaged retinas (Fig. 2e).

**Figure 2.**
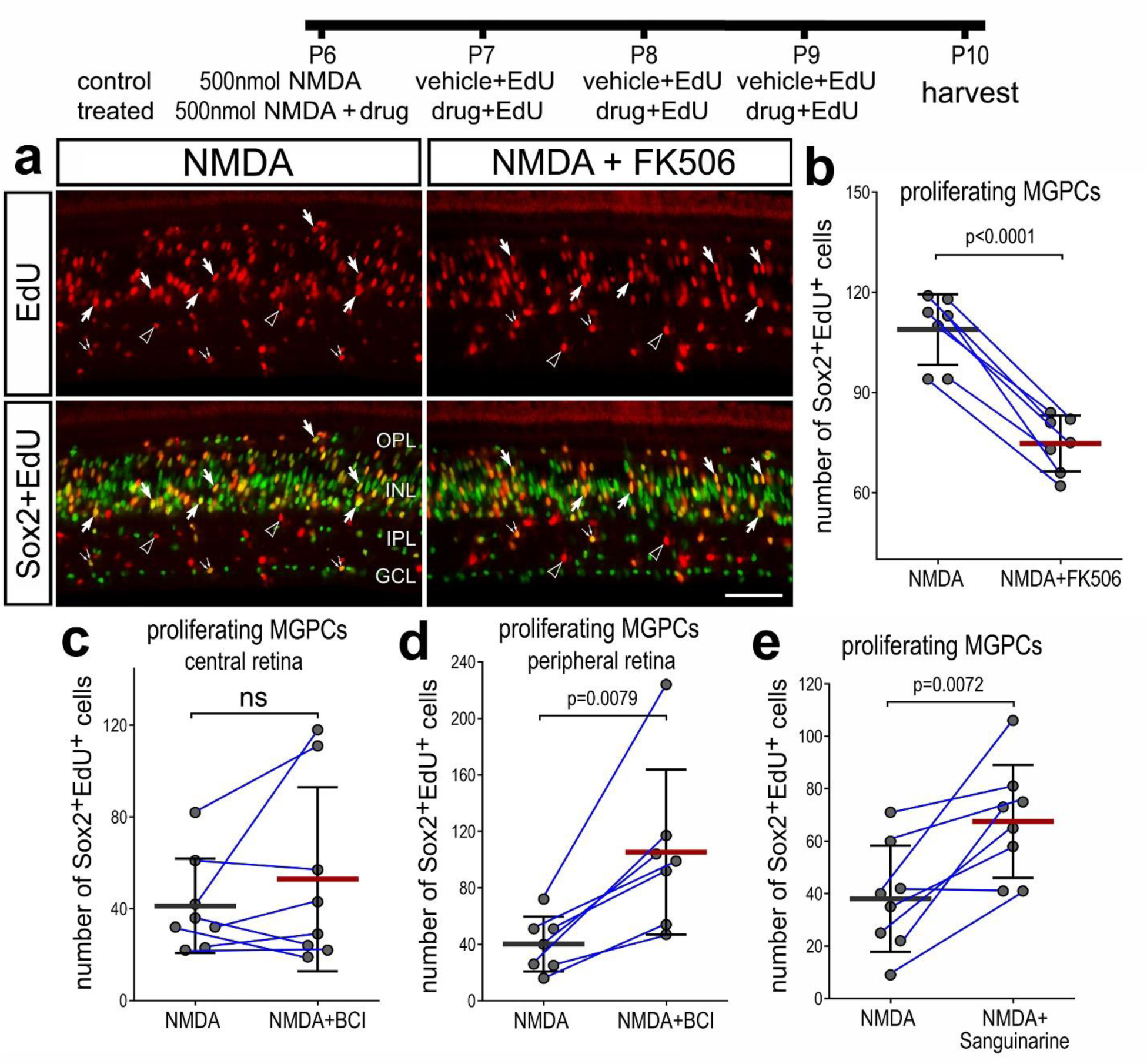
Phosphatase inhibitors influence the proliferation of MGPCs in damaged retinas. Eyes were injected with NMDA ± drug at P6, EdU ± drug at P7, P8 and P9, and retinas harvested at P10. Sections of the retina were labeled for EdU incorporation (red; **a**) and Sox2 (green; **a**). Arrows indicate nuclei of MGPCs, hollow arrow-heads indicate the nuclei of putative microglia, and small double-arrows indicate nuclei of putative NIRG cells. Abbreviations: ONL – outer nuclear layer, INL – inner nuclear layer, IPL – inner plexiform layer, GCL – ganglion cell layer. The calibration bar represents 50 µm (**a**). Histograms (**b-d**) illustrate the mean (bar ± SD), each dot represents one biological replicate and blue lines connect counts from control and treated retinas from one individual. Significance of difference (p-values) was determined by using a paired t-test.

Given the widespread expression of protein phosphatases in retinal cells including amacrine and bipolar neurons which are known to be destroyed by NMDA in the chick retina [36], it is possible that the survival of inner retinal neurons is affected by phosphatase inhibitors. Further, levels of damage to inner retinal neurons are known to influence numbers of proliferating MGPCs [2, 55]. Accordingly, we investigated whether phosphatase inhibitors influenced numbers of dying cells in NMDA damaged retina. We found that there was no significant influence on numbers of dying cells in damaged retinas treated with BCI, sanguinarine or FK506 (Fig. 3a-d).

**Figure 3.**
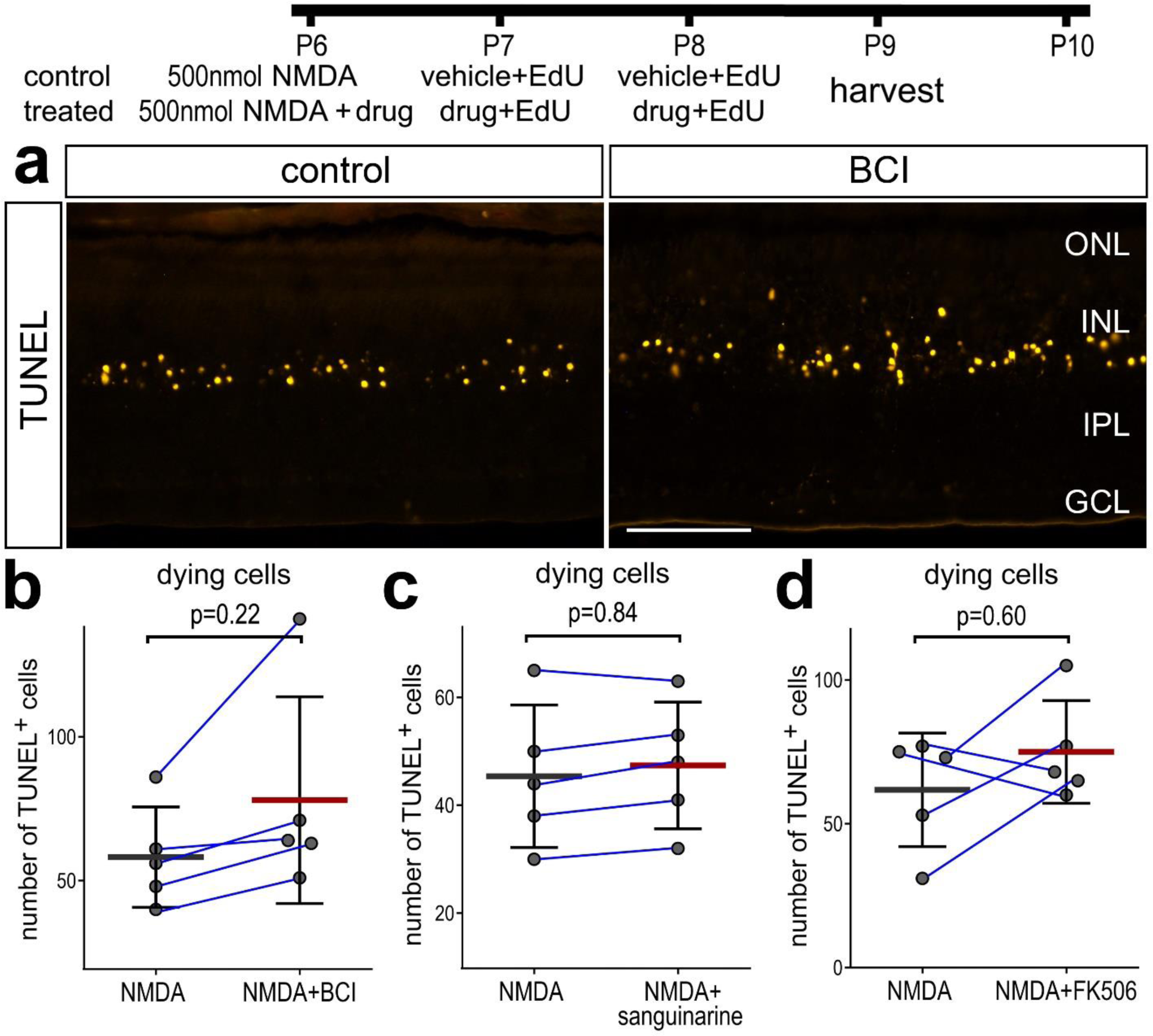
Phosphatase inhibitors do not influence numbers of dying cells in damaged retinas. Eyes were injected with NMDA ± drug at P6, EdU ± drug at P7, P8 and P9, and retinas harvested at P10. Sections of the retina were labeled using TUNEL method (**a**). Significance (p-values) of difference was determined by using a Mann-Whitney U test. Arrows indicate MG and small double arrows indicate amacrine cells. Abbreviations: ONL – outer nuclear layer, INL – inner nuclear layer, IPL – inner plexiform layer, GCL – ganglion cell layer. The calibration bar represents 50 µm (**a**). Histograms (**b-d**) illustrate the mean (bar ± SD), each dot represents one biological replicate and blue lines connect counts from control and treated retinas from one individual. Significance of difference (p-values) was determined by using a Mann-Whitney U-test.

### Inhibition of protein phosphatases and cell signaling

To identify the different cell signaling pathways influenced by BCI or FK506, we probed for readouts of the MAPK pathway (pERK1/2), the Jak/Stat pathway (pStat3) and the mTor-pathway (pS6). In addition, we probed the levels of cFos in MG as readout of the convergence of different cell signaling pathways. We applied BCI or FK506 with NMDA and at 24hrs after NMDA, and harvested retinas at 48hrs after damage when the activation of most cell signaling pathways is subsiding [17–19, 32]. We found that BCI-treatment significantly increased levels of pERK1/2 and cFos in MG in damaged retinas (Fig. 4as,b,e). By comparison, BCI-treatment did not influence levels of pStat3 or pS6 in MG (Fig. 4a,c,d). We found that FK506-treatment significantly decreased levels of pStat3 and pS6 in damaged retinas (Fig. 4f,h,i). By comparison, FK506-treatment did not significantly influence levels of pERK1/2 or cFos in MG (Fig. 4f,g,j).

**Figure 4.**
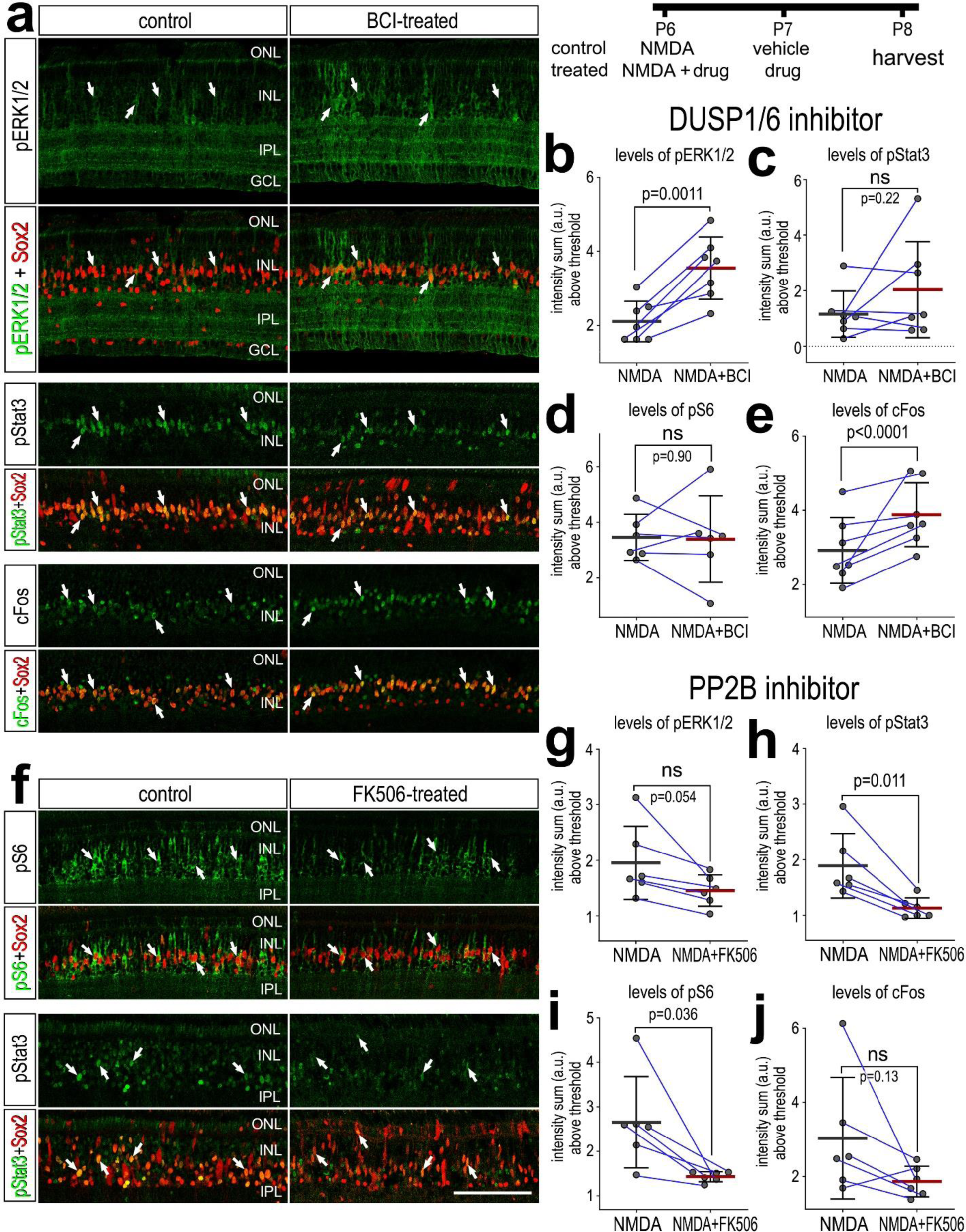
Effects of phosphatase inhibitor on cell signaling in damaged retinas. Eyes were injected with NMDA alone or NMDA + BCI (DUSP1/6 inhibtor) or FK506 (PP2B inhibitor) at P6, vehicle or drug at P7 and retinas harvested at P8. Sections of the retina were labeled with antibodies to pERK1/2 (green; **a**), pStat3 (green; **a,f**), cFos (green; **a**), pS6 (green; **f**) and Sox2 (red; **a,f**). Arrows indicate the cell bodies or nuclei of MG. Abbreviations: ONL – outer nuclear layer, INL – inner nuclear layer, IPL – inner plexiform layer, GCL – ganglion cell layer. The calibration bar represents 50 µm (**a,f**). (**b-e, g-j**) Histograms illustrate the mean (bar ± SD), each dot represents one biological replicate, and blue lines connect counts from control and treated retinas from one individual. Significance of difference (p-values) was determined by using a paired t-test.

### Protein phosphatases in undamaged retinas treated with insulin and FGF2

In the absence of retinal damage, the combination of insulin and FGF2 is known to stimulate the formation of proliferating MGPCs [9, 56]. Thus, we probed for patterns of expression protein phosphatases in scRNA-seq libraries of undamaged retinas treated with 2 or 3 doses of insulin and FGF2 (Fig. 5a,b), as described in detail in previous studies [8, 31, 35, 45]. Treatment with insulin and FGF2 is known to potently impact patterns of gene expression in MG [8, 45, 57]. For example, genes expressed by resting MG, including *GLUL* and *RLBP1*, are robustly downregulated, some genes expressed by activated MG, including *PMP2* and *S100A6*, are robustly upregulated in response to insulin and FGF2, and some genes associated with proliferating MGPCs, including *CDK1* and *TOP2A* are upregulated (Fig. 5c-e). Consistent with patterns of expression seen in other scRNA-seq libraries, feature plots indicated that protein phosphatases were widely expressed by most types of retinal neurons and glia (Fig. 5f). In resting MG, relative levels of protein phosphatases were relatively high and all protein phosphatases, except *PPM1D/E/G*, were significantly downregulated by MG treated with insulin and FGF2 (Fig. 5f,g; Supplemental table 1). Levels of *DUSP1* were significantly increased in MGPCs compared to levels in MG in UMAP clusters treated with insulin and FGF2, whereas levels of other protein phosphatases were not significantly different (Fig. 5g; Supplemental table 1).

**Figure 5.**
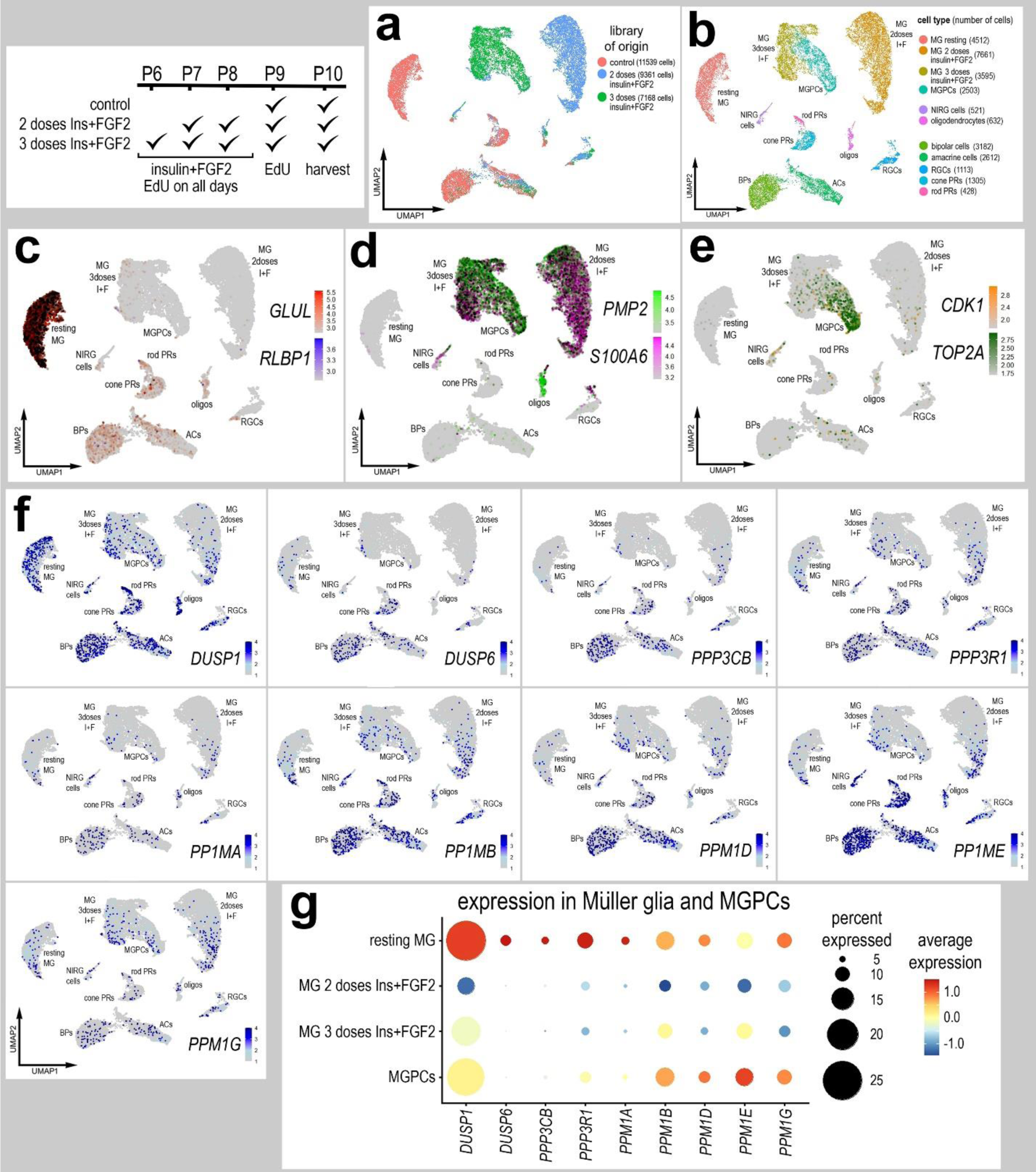
Patterns of expression of protein phosphatases in retinas treated with insulin and FGF2. scRNA-seq was used to identify patterns and levels of expression of protein phosphatases in retinal cells treated with saline, or 2 or 3 doses of insulin and FGF2 retinas (**a**). UMAP clusters of cells were identified based on well-established patterns of gene expression (see Methods; **b**). MG were identified by expression of *RLBP1* and *GLUL* in resting MG (**c**), *S100A6* and *PMP2* in activated MG (**d**), and *CDK1* and *TOP2A* in proliferating MGPCs (**e**). Each dot represents one cell and black dots indicate cells that express 2 or more genes (**c,d,e**). UMAP heatmap plots (**f**) illustrate patterns and levels of expression for phosphatases in retinal neurons and glia. The dot plots in **g** illustrates average expression (heatmap) and percent expressed (dot size) for different phosphatases in MG that are significantly (p<0.0001) up- or downregulated at different times following NMDA-treatment. Significance of difference was determined by using a Wilcox rank sum test with Bonferroni correction.

It is possible that the effects of protein phosphatases upon the formation of MGPCs are impacted by signals produced by damaged neurons and activated microglia/macrophage. Accordingly, we investigated whether phosphatase inhibitors influenced the proliferation of MGPCs in undamaged retinas. Treatment with insulin and FGF2 is known to stimulate the formation of proliferating MGPCs in peripheral regions of the retina in the absence of damage [9]. We found that BCI significantly increased numbers of proliferating MGPCs, but did not significantly influence the percent EdU-labeled MGPCs that expressed Sox2 (Fig. 6a,b,c). Interestingly, sanguinarine had no effect upon the proliferating of MGPCs in retinas treated with insulin and FGF2 (Fig. 6d). Interestingly, FK506 stimulated the proliferation of MGPCs in retinas treated with insulin and FGF2 (Fig. 6e), the opposite effect of this inhibitor on MGPC proliferation in NMDA-damaged retinas. This finding suggests that there are significant differences in MG treated with insulin+FGF2 and NMDA that may underlie the different effects of FK506 on the proliferation of MGPCs in undamaged and damaged retinas.

**Figure 6.**
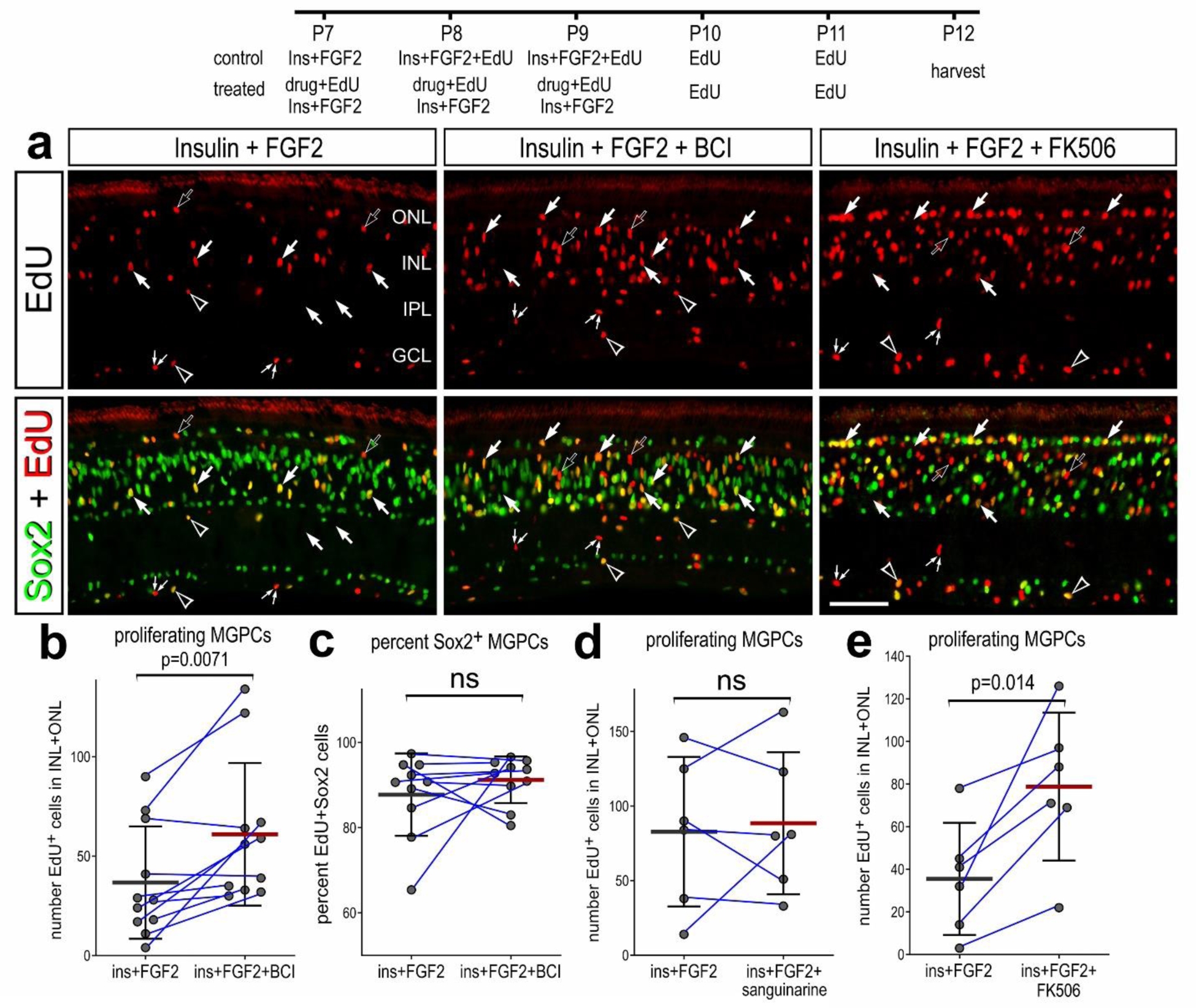
Phosphatase inhibitors influence the proliferation of MGPCs in undamaged retinas treated with insulin and FGF2. Eyes were injected with insulin and FGF2 ± drug at P7, P8 and P9, EdU at P10 and P11 and retinas harvested at P12. Sections of the retina were labeled for EdU incorporation (red; **a**) and Sox2 (green; **a**). Arrows indicate nuclei of Sox2+/EdU+ MGPCs, hollow arrows indicate Sox2-/EdU+ MGPCs, hollow arrow-heads indicate the nuclei of putative NIRG cells, and small double-arrows indicate nuclei of putative microglia. Abbreviations: ONL – outer nuclear layer, INL – inner nuclear layer, IPL – inner plexiform layer, GCL – ganglion cell layer. The calibration bar represents 50 µm (**a**). (**b-d**) Histograms illustrate the mean (bar ± SD), each dot represents one biological replicate, and blue lines connect counts from control and treated retinas from one individual. Significance of difference (p-values) was determined by using a paired t-test.

To better understand the different effects of FK506 on MGPCs, we probed for differentially expressed genes (DEGs) in MG in a large aggregated scRNA-seq database of retinal cells that included 82,765 MG from retinas treated saline, NMDA, insulin+FGF2 and NMDA+insulin+FGF2 (Fig. 7a,b). We generated and analyzed these scRNA-seq libraries in previous studies [8, 45, 58]. UMAP ordering of cells revealed distinct clusters of neurons and glia based on patterns of expression of cell-distinguishing markers, with distinct clusters of MG based on different treatments (Fig. 7a-c). We identified the up- and downregulated DEGs in MG from retinas treated with 2 doses of insulin+FGF2 and 24hr after NMDA, when MG are known to undergo the process of reprogramming before re-entering the cell cycle [8, 59, 60]. We identified numerous upregulated DEGs in MG from retinas treated with saline vs 24hr NMDA or 2doses insulin+FGF2 and for MG from retinas treated with 2 doses insulin+FGF2 vs 24hr NMDA (Fig. 7d,e). Compared to MG treated with 2 doses of insulin+FGF2, MG from retinas 24hrs after NMDA significantly upregulated *DUSP1, DUSP6* and *PPP3CA*, along with different transcription factors (TFs), components of different cell signaling pathways and secreted factors (Fig. 7d,e). We next performed Gene Ontology (GO) enrichment analysis on upregulated DEGs in MG from retinas 24hr after NMDA compared to MG treated with 2 doses of insulin+FGF2 (Fig. 7f). We identified many different gene modules related to cell signaling pathways that have been implicated in regulating the formation of MGPCs included Wnt- [22], Hippo- [61, 62], Jak/Stat [19] , TNF- [63, 64], Hedgehog [24, 25], Notch- [26, 27, 65, 66], TGFβ- [29, 34] and mTO [32]. In addition, we identified modules of upregulated genes associated with neurogenesis, cell cycle and regulation of protein phosphorylation (Fig. 7f). Importantly, we identified upregulated DEGs implicating gene modules for protein phosphatase activity (Fig. 7f), suggesting that MG in damaged retinas expressed more phosphatase inhibitor targets compared to MG in retinas treated with insulin and FGF2.

**Figure 7.**
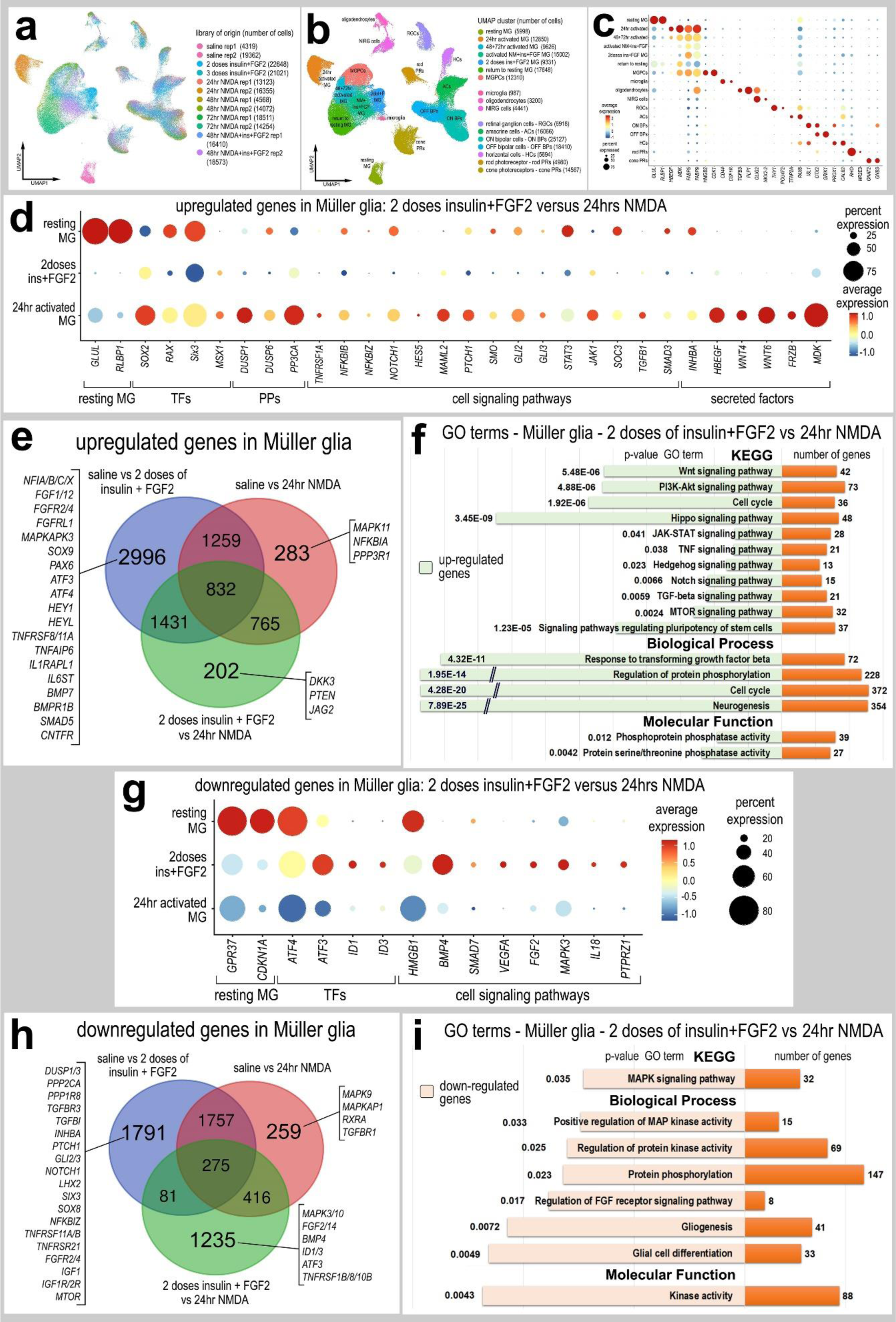
Comparison of protein phosphatase expression in MG from retinas treated with NMDA versus insulin+FGF2. scRNA-seq was used to compare patterns of expression of protein phosphatases in MG in undamaged retinas treated with 2 doses of insulin+FGF2 and damaged retinas treated with NMDA. UMAP plots illustrate a total of 183,216 cells from different treatments and replicates (**a**). MG formed distinct UMAP clusters that correlated to different treatments and MGPCs based on progression through the cell cycle (**a-c**). MG were bioinformatically isolated from scRNA-seq libraries of saline-treated retinas (5887 MG), retinas at 24 after NMDA treatment (12,667 MG), and retinas treated with 2 doses of insulin and FGF2 (9047 MG). Lists of DEGs were generated (Supplemental tables 2-4) for MG from retinas treated with saline vs 24hr NMDA, saline vs 2 doses of insulin+FGF2, and 2 doses of insulin+FGF2 vs 24hr NMDA. Numbers of up- and down-regulated DEGs are plotted in a Venn diagrams (**e,h**). Genes of interest unique to each treatment are listed. Dot plots illustrate the percentage of expressing MG (size) and significant (p<0.0001) changes, upregulated (**d**) and downregulated (**g**), in expression levels (heatmap) for genes in MG from retinas treated with 2 doses of insulin+FGF2 vs 24hr NMDA. Significance of difference was determined by using a Wilcox rank sum test with Bonferroni correction (Supplemental tables 2-4). GO enrichment analysis was performed for lists of DEGs in MG. Gene modules for upregulated (green; **f**) and downregulated (peach; **i**) genes were grouped by GO category with P values and numbers of genes for each category.

We next identified numerous downregulated DEGs in MG from retinas treated with saline vs 24hr NMDA or 2 doses insulin+FGF2 and for MG from retinas treated with 2 doses insulin+FGF2 vs 24hr NMDA (Fig. 7g,h). Compared to MG treated with 2doses of insulin+FGF2, MG from retinas 24hrs after NMDA significantly downregulated different transcription factors (TFs) associated with reactivity and glial specification and components of different cell signaling pathways (Fig. 7g,h). We performed GO enrichment analysis on downregulated DEGs in MG from retinas treated 2 doses insulin+FGF2 vs 24hr after NMDA (Fig. 7i). We identified many different gene modules related to MAPK signaling, protein kinase activity and glial cell development and differentiation (Fig. 7i).

### Expression of protein phosphatases in MG in retinas missing microglia

We next investigated whether microglia and macrophage influenced the expression of phosphatases in MG in healthy and damaged retinas. Given that we do not distinguish between microglia and macrophage, hereafter we use the term microglia to collectively refer to microglia and macrophage. Retinal microglia are known to influence the ability of MG to become proliferating MGPCs [56, 67]. Accordingly, we probed scRNA-seq of normal and damaged chick retinas with the microglia intact or ablated by clodronate-liposomes, as originally described in detail in a recent study [68]. More than 95% of the microglia are ablated with 3 days of treatment with clodronate-liposome [39, 56]. UMAP plots of aggregated libraries (43,566 total cells) contained distinct clusters of different cell types, with neuronal cells clustered together independent of treatment whereas MG forming distinct clusters that correlated to treatment (Fig. 8a). Clusters of retinal cells were identified based on cell-distinguishing genes (see Methods). Resting MG were identified based on high levels of expression for *GLUL, RLBP1* and *GPR37L1*, and activated MG were identified based on upregulation of *PMP2, TGFB2* and *MDK* (Fig. 8b). MG were bioinformatically isolated and re-ordered in a UMAP plot. Approximately 1000 MG from each treatment group formed distinct clusters of cells that were segregated into resting and activated cells from different treatments (Fig. 8c,d). In undamaged retinas, resting MG expressed relatively low levels of *DUSP1, DUSP6, PPP3CB, PPP3R1* and *PPM1B/D/E/G,* whereas PPM1A was relatively high in resting MG (Fig. 8e,f). Ablation of microglia from undamaged retinas did not result in significant changes in expression of protein phosphatases (Fig. 8e,f; Supplemental table 1). Consistent with data from other scRNA-seq libraries, levels of *DUSP1, DUSP6, PPP3CB, PPP3R1* and *PPM1A/B/E/G* are significantly increased at 24 hrs after NMDA-treatment (Fig. 8e,f). In damaged retinas missing microglia, levels of *DUSP1* were significantly reduced whereas levels of other phosphatases were not significantly different (Fig. 8e,f; Supplemental table 1).

**Figure 8.**
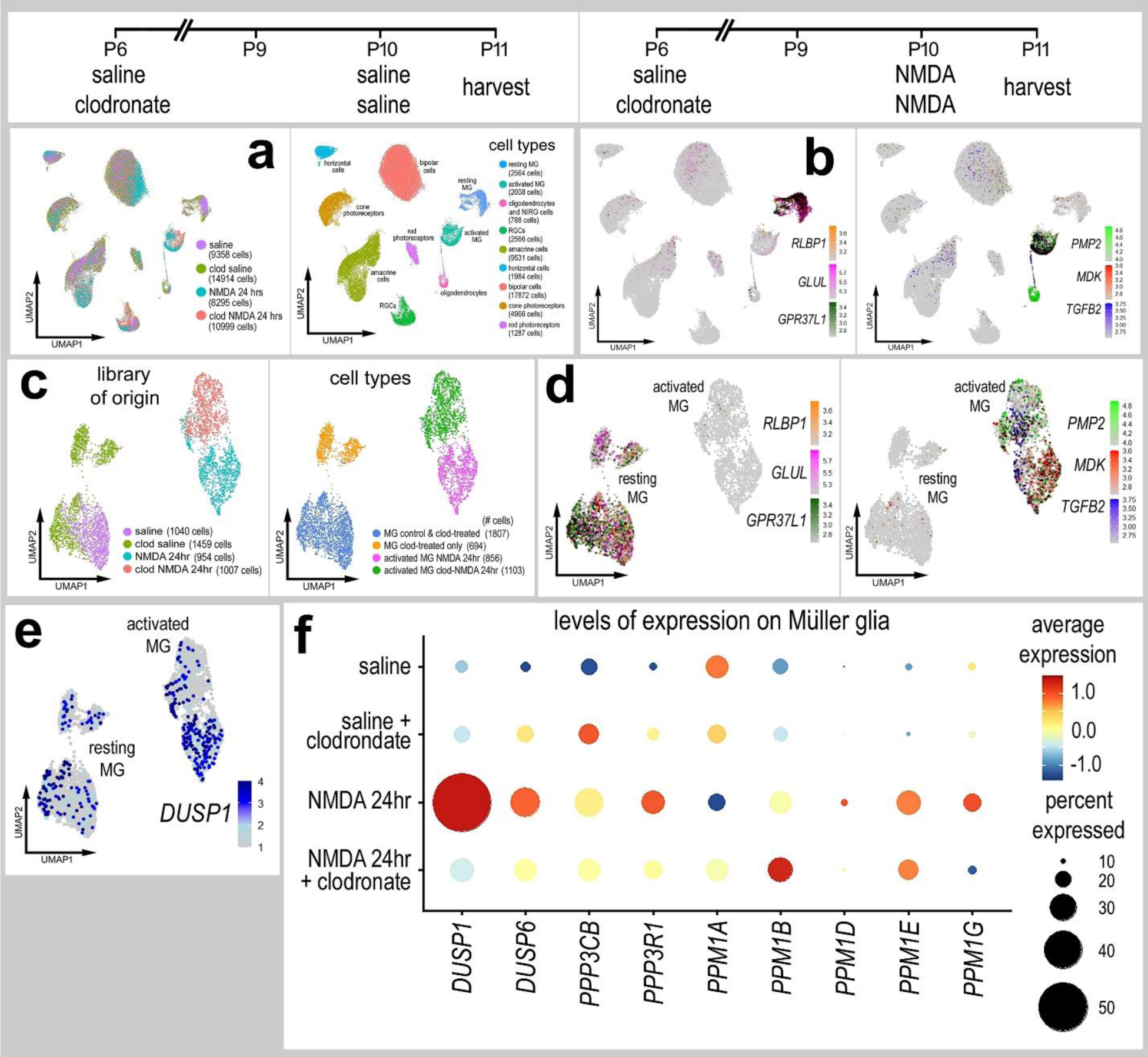
Expression of protein phosphatases in normal and damaged retinas with and without microglia. Retinas were obtained from eyes injected with saline or clodronate liposomes at P6, saline or NMDA at P10, and tissues harvested at 24hrs after the last injection. UMAP ordering of cells is displayed for libraries of origin and distinct clusters of cells (**a**). In UMAP heatmap plots, resting MG were identified based on elevated expression of *RLBP1, GLUL* and *GPR37L1* (**b,d**), and activated MG were identified based on elevated expression of *PMP2, MDK* and *TGFB2* (**b,d**). MG were bioinformatically isolated and re-embedded in UMAP plots (**c,d**). UMAP heatmap plots (**e**) illustrate patterns and levels of expression for protein phosphatases in MG. The dot plot in **f** illustrates average expression (heatmap) and percent expressed (dot size) for different phosphatases in MG that are significantly (p<0.0001) up- or downregulated. Significance of difference was determined by using a Wilcox rank sum test with Bonferroni correction.

## Discussion

The protein phosphatases that were represented in our scRNA-seq libraries are rapidly upregulated and then downregulated in MG following NMDA-induced damage. By comparison, these protein phosphatases are downregulated in MG in undamaged retinas in response to treatment with insulin+FGF2. Following retinal damage or treatment with insulin+FGF2, numerous different kinase-dependent cell signaling pathways are known to be rapidly activated in MG in the chick retina. These pathways include MAPK [17, 18], Jak/Stat [19], BMP/SMAD1/5/9 [29], Wnt/β-catenin [22] and NFkB [35]. Thus, the rapid downregulation of protein phosphatases in MG likely facilitates the activation of many different kinase-dependent signaling cascades in response retinal damage or insulin and FGF2 in the absence. This may explain some of the differences observed in the formation of MGPCs with different protein phosphatase inhibitors in NMDA-damaged retinas and retinas treated with insulin and FGF2. Thus, it is expected that different protein phosphatase act to fine-tune the kinase-driven responses of MG to damage or growth factors.

### Specificity of effects and different families of Protein Phosphatases

DUSP1 and DUSP6 are known to dephosphorylate different MAPKs [69, 70], and dephosphorylation of MAPKs is expected to suppress the activity of this pathway. Thus, since activation of MAPK is known to stimulate the formation of proliferating MGPCs in the chick retina [17, 18], inhibition of DUSP1/6 is expected to facilitate the formation of MGPCs. Indeed, our findings indicate that DUSP1/6 inhibitor increased levels of pERK1/2 and cFos, and resulted in increased numbers of proliferating MGPCs, but only in peripheral regions of NMDA-damaged retinas and retinas treated with insulin and FGF2. By comparison, FK506 is known to inhibit the PP2B family of phosphatases, which includes PPP3CB (Protein Phosphatase 3 Catalytic Subunit Beta) which is best known to act at NFAT (Nuclear Factor of Activated T cells), p53 (Tumor Protein p53), ELK1 (ETS Transcription Factor ELK1) [71–73]. Calcineurin is also known act at different components of the NFkB pathway to result in suppression of NFkB-signaling [50, 74]. NFkB-signaling is known to suppress the formation of proliferating MGPCs in the chick retina [35]. Thus, it is possible that the inhibition of calcineurin secondarily facilitated NFkB-signaling in MG and thereby decreased numbers of proliferating MGPCs in damaged retinas. Further, we find that PP2B inhibitor decreased activation of Jak/Stat and mTor pathways in MG which have been shown promote the formation of proliferating MGPCs [19, 32]. The decreased signaling through Jak/Stat and mTor pathways in MG in damaged retinas treated with PP2B inhibitor is likely secondary to activation of inhibitory pathways that are inactivated by PP2B phosphatases.

The actions of PP2C inhibitor (sanguinarine) are more difficult to interpret. It is possible that sanguinarine may act at PPM1E which inhibits PAK, CAMK2/4 and p38 MAPK [75, 76]. Alternatively, sanguinarine may act at PPM1B which is known to dephosphorylate cyclin-dependent kinases (CDKs), causing cell growth arrest or cell death [77]. CDK1 and CDK6 are highly expressed by MGPCs (Fig. 4e; not shown) and inactivation of CDK’s is expected to suppress the proliferation of MGPCs, which was not observed for treatment of damaged retinas with sanguinarine. In addition, PPM1B dephosphorylates IKKβ to terminate TNFα-induced NFkB activation [78], and inhibition of NFkB is known to stimulate the proliferation of MGPCs [35]. Sanguinarine is also known to inhibit PPM1A which is known to deactivate pSMAD2/3 [52, 79]; SMAD3-signaling is known to suppress the formation of MGPCs in the chick retina [29]. Thus, inhibition of PPM1A by sanguinarine might be expected to increase MGPC proliferation by suppressing SMAD3- and NFkB-signaling. Further studies are required to determine the precise mechanisms of action of sanguinarine in the formation of MGPCs in the chick retina.

### Effects in peripheral retina

We found that DUSP1/6 inhibitor stimulated the proliferation of MGPCs in peripheral regions of retinas, but not central retina. This may have resulted from inherent differences between MG found in central and peripheral regions of the retina. For example, as the chicks age, the MG that respond to damage by proliferating as MGPCs in regions of retina that progressively become confined to narrower regions of the peripheral retina [59]. Similarly, the effects of insulin+FGF2 on MGPC formation are most prominent in peripheral regions of the retina, whereas MG in central regions of the retina do not reprogram [9]. It is possible that some of the differences between MG in central and peripheral regions of retina are imparted by Pax2, which is expressed by MG in central retinas and not in peripheral regions retina [44].

### Amplitude of effects

Potent inhibition of MGPCs formation includes treatments such as ablation of microglia [56], inhibition of mTor [32], inhibition of Notch [26, 27], activation of glucocorticoid signaling [33], inhibition of FGFR/MAPK [17, 18], inhibition of fatty acid binding proteins [8, 45], and inhibition of S-adenosyl homocysteine hydrolase [48]. Pathways that have relatively subtle effects on numbers of proliferating MGPCs in damaged retinas include Hedgehog [25], Jak/Stat [19], retinoic acid [28], endocannabinoid signaling [30], activation of BMP/Smad1/5/8, inhibition of TGFβ/Smad3 [29], inhibition of MMP [80], midkine [57], and NFkB-signaling [35]. However, NFkB has time and context specific effects upon MG; NFkB-signaling is required for the rapid activation of MG (early first steps in reprogramming), but thereafter suppresses the proliferative and neurogenic capacity of MGPCs [35]. Nevertheless, different phosphatase inhibitors had relatively modest effects on MGPC proliferation, suggesting that the actions of phosphatases “fine tune” the kinase-dependent cascades. It is also possible that the phosphatase inhibitors are influencing different pathways that simultaneously stimulate and suppress MG reprogramming to result in relatively modest effects on MGPC proliferation.

### Opposite effects of PP2B inhibitor on MGPCs in damaged and undamaged retinas

In damaged retinas, we found that PP2B inhibitor, FK506, decreased the proliferation of MGPCs by about 30%. By contrast, in undamaged retinas treated with insulin and FGF2, PP2B inhibitor increased the proliferation of MGPCs by more than 100%. We have identified different factors that influence the proliferation of MGPCs in damaged retinas that have no effect on MGPCs in undamaged retinas, including ID transcription factors [81], midkine [57], and HBEGF [47]. The current findings regarding the effects of PP2B inhibitor on MGPCs proliferation are the first to show opposite effects in damaged and undamaged retinas. To better understand the differential effects of PP2B inhibitor, we identified differential gene expression in MG in retinas at 24hrs after NMDA and 2 doses of insulin and FGF2 when these cells undergoing the process of reprogramming prior to going through the cell cycle. We identified numerous upregulated DEGs in MG in NMDA damaged retinas, compared to MG in retinas treated with insulin and FGF2. The upregulated DEGs are associated with gene modules associated with protein phosphatase activity and numerous cell signaling pathways that are known to drive the formation of MGPCs. Our findings suggest that the increased phosphatase-related gene modules include FK506-sensitive phosphates that inactivate pathways that suppress the formation of proliferating MGPCs in damaged retinas, which may include TNF/NFkB- [35] and TGFβ/Smad2/3-signaling [29]. However, in undamaged retinas TNF/NFkB- and TGFβ/Smad-signaling are known to suppress the formation of MGPCs [29, 35]. Our findings suggest that there is less activation of TNF/NFkB- and TGFβ/Smad-signaling in undamaged insulin/FGF2-treated retina compared to levels seen in damaged retinas. However, this does not explain the opposite effects of PP2B inhibitor on MGPC proliferation. In retinas treated with insulin and FGF2 we identified more DEGs associated with FGF/MAPK-signaling and generic kinase activity. FGF/MAPK-signaling is known to drive the formation of proliferating MGPCs [17, 18]. These pathways may include an FK506-sensitive phosphatases that suppresses the formation of MGPCs in undamaged retinas.

## Conclusions

Our findings indicate that different classes of protein phosphatases are dynamically expressed by MG and MGPCs in damaged retinas or retinas treated with insulin and FGF2 in the absence of damage. Inhibition of different classes of protein phosphatases has differential effects on the formation of proliferating MGPCs in damaged retinas and undamaged retinas treated with insulin and FGF2. Our findings suggest that PP2B family of phosphatases have context-specific influences of the formation of MGPCs that differ between damaged and undamaged retinas. Collectively, these findings are consistent with the notion that different kinase-dependent cell signaling cascades control the reprogramming of MG into proliferating MGPCs, and protein phosphatases fine-tune this process.

## Data availability

Cell Ranger output files for Gene-Cell matrices for scRNA-seq data for libraries from saline and NMDA-treated retinas are available through GitHub: https://github.com/jiewwwang/Singlecell-retinal-regeneration or Sharepoint: [chick embryonic retina scRNA-seq Cell Ranger outs](https://osumc.sharepoint.com/sites/Links/FischerLab/Forms/AllItems.aspx?id=%2Fsites%2FLinks%2FFischerLab%2Fchick%20embryonic%20retina%20scRNA%2Dseq%20Cell%20Ranger%20outs&p=true&ga=1) (https://osumc.sharepoint.com/:f:/s/Links/Eoto-Qg2uuxDn1bHWMM6gdkBTft4S_YSBjQJResxY-qehA?e=mdaPlg) and [chick retina scRNA-seq Cell Ranger output files](https://osumc.sharepoint.com/sites/Links/FischerLab/Forms/AllItems.aspx?id=%2Fsites%2FLinks%2FFischerLab%2Fchick%20retina%20scRNA%2Dseq%20Cell%20Ranger%20output%20files&p=true&ga=1) (https://osumc.sharepoint.com/:f:/s/Links/Eoto-Qg2uuxDn1bHWMM6gdkBTft4S_YSBjQJResxY-qehA?e=mdaPlg). scRNA-seq datasets are deposited in GEO (GSE135406, GSE242796) and Gene-Cell matrices for scRNA-seq data for libraries from saline and NMDA-treated retinas are available through NCBI (GSM7770646, GSM7770647, GSM7770648, GSM7770649).

## Acknowledgements

This work was supported by RO1 EY022030-09 and RO1 EY032141-03 (AJF).

---

**Supplemental Table 1:** Statistics for genes listed in Dot Plots in Figures 1, 5 and 8.

**Supplemental Table 2:** Differentially Expressed Genes (DEGs) in Müller glia in retinas treated with saline (pct.1) vs 24hrs after NMDA-treatment (pct.2). Negative avg log2FC represents an increase and positive avg log2FC represents a decrease.

**Supplemental Table 2:** Differentially Expressed Genes (DEGs) in Müller glia in retinas treated with saline (pct.1) vs 2 doses of insulin+FGF2 (pct.2). Negative avg log2FC represents an increase and positive avg log2FC represents a decrease.

**Supplemental Table 4:** Differentially Expressed Genes (DEGs) in Müller glia in retinas treated with 2 doses of insulin+FGF2 (pct.1) vs 24hrs after NMDA-treatment (pct.2). Negative avg log2FC represents an increase and positive avg log2FC represents a decrease.

