## Supplemental Table 4 for "Protein phosphatases regulate the formation of Müller glia-derived progenitor cells in the chick retina"

**Statistics for dotplot figure 1n**

```
FindMarkers(early_MG, ident.1 = "saline", ident.2 = c("NM3hrs", "NM12hrs", "NM48hrs"), features = c("DUSP1", "DUSP6", "PPP3CB", "PPP3R1", "PPM1A", "PPM1B", "PPM1D", "PPM1E", "PPM1G"), group.by = "orig.ident", min.pct = 0.005, logfc.threshold = 0.02)
```

| | p_val | avg_log2FC | pct.1 | pct.2 | p_val_adj |
|---|---|---|---|---|---|
| DUSP6 | 5.371206e-114 | -0.45180127 | 0.053 | 0.375 | 1.054905e-109 |
| PPM1E | 3.571155e-91 | -0.53839369 | 0.279 | 0.604 | 7.013748e-87 |
| DUSP1 | 1.040336e-52 | -0.56949698 | 0.317 | 0.580 | 2.043219e-48 |
| PPM1G | 1.069230e-39 | -0.22707246 | 0.245 | 0.480 | 2.099967e-35 |
| PPP3CB | 1.372310e-30 | -0.12768066 | 0.263 | 0.475 | 2.695216e-26 |
| PPP3R1 | 2.188437e-20 | -0.11577262 | 0.375 | 0.573 | 4.298090e-16 |
| PPM1B | 1.536903e-17 | -0.10817286 | 0.347 | 0.532 | 3.018477e-13 |
| PPM1D | 4.537538e-13 | 0.02814839 | 0.179 | 0.303 | 8.911724e-09 |
| PPM1A | 1.520403e-09 | 0.06635710 | 0.222 | 0.343 | 2.986071e-05 |

```
FindMarkers(early_MG, ident.1 = "saline", ident.2 = "NM3hrs", features = c("DUSP1", "DUSP6", "PPP3CB", "PPP3R1", "PPM1A", "PPM1B", "PPM1D", "PPM1E", "PPM1G"), group.by = "orig.ident", min.pct = 0.005, logfc.threshold = 0.02)
```

| | p_val | avg_log2FC | pct.1 | pct.2 | p_val_adj |
|---|---|---|---|---|---|
| DUSP6 | 1.184720e-114 | -0.65106680 | 0.053 | 0.452 | 2.326790e-110 |
| PPM1E | 6.298443e-113 | -0.99953037 | 0.279 | 0.728 | 1.237014e-108 |
| PPP3R1 | 7.576690e-33 | -0.40855398 | 0.375 | 0.685 | 1.488062e-28 |
| PPM1G | 7.820290e-30 | -0.19835881 | 0.245 | 0.526 | 1.535905e-25 |
| PPP3CB | 2.732089e-23 | -0.19565867 | 0.263 | 0.514 | 5.365822e-19 |
| PPM1A | 4.263695e-18 | -0.08282910 | 0.222 | 0.447 | 8.373896e-14 |
| PPM1D | 1.200563e-17 | -0.07589580 | 0.179 | 0.375 | 2.357906e-13 |
| PPM1B | 5.498413e-10 | -0.14116835 | 0.347 | 0.551 | 1.079888e-05 |
| DUSP1 | 1.540236e-08 | -0.09944786 | 0.317 | 0.495 | 3.025024e-04 |

```
FindMarkers(early_MG, ident.1 = "saline", ident.2 = "NM12hrs", features = c("DUSP1", "DUSP6", "PPP3CB", "PPP3R1", "PPM1A", "PPM1B", "PPM1D", "PPM1E", "PPM1G"), group.by = "orig.ident", min.pct = 0.005, logfc.threshold = 0.02)
```

| | p_val | avg_log2FC | pct.1 | pct.2 | p_val_adj |
|---|---|---|---|---|---|
| DUSP6 | 2.912278e-142 | -0.54779868 | 0.053 | 0.467 | 5.719714e-138 |
| PPM1E | 2.330731e-94 | -0.48371960 | 0.279 | 0.677 | 4.577555e-90 |
| DUSP1 | 2.300063e-93 | -0.85730125 | 0.317 | 0.705 | 4.517323e-89 |
| PPM1G | 2.186273e-48 | -0.28685678 | 0.245 | 0.539 | 4.293840e-44 |
| PPP3CB | 8.551572e-41 | -0.12846678 | 0.263 | 0.542 | 1.679529e-36 |
| PPM1B | 1.458251e-24 | -0.09244339 | 0.347 | 0.586 | 2.864004e-20 |
| PPP3R1 | 1.339483e-17 | -0.02527234 | 0.375 | 0.597 | 2.630746e-13 |
| PPM1D | 1.095109e-16 | 0.04792895 | 0.179 | 0.340 | 2.150794e-12 |
| PPM1A | 2.437147e-10 | 0.07329386 | 0.222 | 0.363 | 4.786557e-06 |

```
FindMarkers(early_MG, ident.1 = "saline", ident.2 = "NM48hrs", features = c("DUSP1", "DUSP6", "PPP3CB", "PPP3R1", "PPM1A", "PPM1B", "PPM1D", "PPM1E", "PPM1G"), group.by = "orig.ident", min.pct = 0.005, logfc.threshold = 0.02)
```

| | p_val | avg_log2FC | pct.1 | pct.2 | p_val_adj |
|---|---|---|---|---|---|
| DUSP6 | 4.369193e-13 | -0.03436052 | 0.053 | 0.145 | 8.581094e-09 |
| DUSP1 | 7.933132e-07 | -0.34084934 | 0.317 | 0.433 | 1.558067e-02 |
| PPM1G | 3.495407e-04 | -0.14164903 | 0.245 | 0.336 | 1.000000e+00 |
| PPM1E | 1.126530e-03 | -0.09599445 | 0.279 | 0.367 | 1.000000e+00 |
| PPP3CB | 6.820175e-02 | -0.06395559 | 0.263 | 0.323 | 1.000000e+00 |
| PPM1B | 7.482898e-02 | -0.10658464 | 0.347 | 0.418 | 1.000000e+00 |
| PPM1A | 1.445115e-01 | 0.19754638 | 0.222 | 0.218 | 1.000000e+00 |
| PPM1D | 3.007826e-01 | 0.08918530 | 0.179 | 0.176 | 1.000000e+00 |

```
FindMarkers(early_MG, ident.1 = "NM3hrs", ident.2 = "NM12hrs", features = c("DUSP1", "DUSP6", "PPP3CB", "PPP3R1", "PPM1A", "PPM1B", "PPM1D", "PPM1E", "PPM1G"), group.by = "orig.ident", min.pct = 0.005, logfc.threshold = 0.02)
```

| | p_val | avg_log2FC | pct.1 | pct.2 | p_val_adj |
|---|---|---|---|---|---|

```
DUSP1  5.438830e-38 -0.75785339 0.495 0.705 1.068186e-33
PPM1E  4.277554e-22  0.51581077 0.728 0.677 8.401117e-18
PPP3R1 1.031266e-08  0.38328165 0.685 0.597 2.025406e-04
PPM1A  6.995902e-04  0.15612295 0.447 0.363 1.000000e+00
PPM1B  4.705885e-03  0.04872496 0.551 0.586 1.000000e+00
PPP3CB 9.608624e-02  0.06719189 0.514 0.542 1.000000e+00
PPM1D  1.255734e-01  0.12382475 0.375 0.340 1.000000e+00
PPM1G  1.851401e-01 -0.08849797 0.526 0.539 1.000000e+00
DUSP6  6.700840e-01  0.10326812 0.452 0.467 1.000000e+00
```

```
FindMarkers(early_MG, ident.1 = "NM12hrs", ident.2 = "NM48hrs", features = c(
"DUSP1", "DUSP6", "PPP3CB", "PPP3R1", "PPM1A", "PPM1B", "PPM1D", "PPM1E", "PP
M1G"), group.by = "orig.ident", min.pct = 0.005, logfc.threshold = 0.02)
              p_val avg_log2FC pct.1 pct.2    p_val_adj
DUSP6  7.307268e-57 0.51343816 0.467 0.145 1.435147e-52
PPM1E  5.727610e-50 0.38772515 0.677 0.367 1.124903e-45
DUSP1  7.994397e-37 0.51645190 0.705 0.433 1.570100e-32
PPP3CB 5.201983e-24 0.06451119 0.542 0.323 1.021669e-19
PPM1G  1.131276e-20 0.14520776 0.539 0.336 2.221827e-16
PPM1D  3.931422e-17 0.04125634 0.340 0.176 7.721312e-13
PPM1A  1.160373e-13 0.12425252 0.363 0.218 2.278973e-09
PPP3R1 3.628250e-12 0.04492779 0.597 0.433 7.125882e-08
```

**Statistics for dotplot figure 5g**

```
FindMarkers(insulin_FGF2_AGG, ident.1 = "resting MG", ident.2 = "MG 2do I+F",
features = c("DUSP1", "DUSP6", "PPP3CB", "PPP3R1", "PPM1A", "PPM1B", "PPM1D",
"PPM1E", "PPM1G"), min.pct = 0.005, logfc.threshold = 0.02)
              p_val avg_log2FC pct.1 pct.2    p_val_adj
DUSP1  2.452758e-94 0.87368445 0.252 0.117 3.644553e-90
DUSP6  1.368140e-69 0.38585988 0.074 0.013 2.032919e-65
PPP3R1 6.257829e-17 0.22042113 0.107 0.065 9.298508e-13
PPM1B  3.415854e-14 0.20066872 0.121 0.080 5.075617e-10
PPP3CB 3.701598e-12 0.13965857 0.055 0.030 5.500205e-08
PPM1A  1.476602e-11 0.15079573 0.060 0.034 2.194082e-07
PPM1D  6.038305e-04 0.08602030 0.079 0.064 1.000000e+00
PPM1G  4.130049e-03 0.08393707 0.100 0.086 1.000000e+00
PPM1E  4.570456e-03 0.05010188 0.110 0.094 1.000000e+00
```

```
FindMarkers(insulin_FGF2_AGG, ident.1 = "resting MG", ident.2 = "MG 3do I+F",
features = c("DUSP1", "DUSP6", "PPP3CB", "PPP3R1", "PPM1A", "PPM1B", "PPM1D",
"PPM1E", "PPM1G"), min.pct = 0.005, logfc.threshold = 0.02)
              p_val avg_log2FC pct.1 pct.2    p_val_adj
DUSP6  3.066857e-28 0.32051911 0.074 0.020 4.557043e-24
PPP3R1 8.895700e-16 0.24783921 0.107 0.057 1.321812e-11
DUSP1  3.160732e-14 0.42550411 0.252 0.187 4.696532e-10
PPP3CB 1.877340e-13 0.17797616 0.055 0.023 2.789539e-09
PPM1A  5.432182e-09 0.14606615 0.060 0.032 8.071679e-05
PPM1D  5.224742e-05 0.08572252 0.079 0.056 7.763445e-01
PPM1G  6.508550e-04 0.10186121 0.100 0.078 1.000000e+00
PPM1B  4.440572e-03 0.04957836 0.121 0.100 1.000000e+00
```

```
FindMarkers(insulin_FGF2_AGG, ident.1 = "resting MG", ident.2 = "MGPCs", feat
ures = c("DUSP1", "DUSP6", "PPP3CB", "PPP3R1", "PPM1A", "PPM1B", "PPM1D", "PP
M1E", "PPM1G"), min.pct = 0.005, logfc.threshold = 0.02)
                p_val  avg_log2FC pct.1 pct.2    p_val_adj
DUSP6  5.888801e-23  0.35017020 0.074 0.018 8.750170e-19
PPP3CB 5.103169e-06  0.13571988 0.055 0.031 7.582799e-02
PPP3R1 3.024334e-05  0.15788130 0.107 0.077 4.493858e-01
PPM1A  1.631852e-03  0.09630454 0.060 0.042 1.000000e+00
```

```
DUSP1  1.315107e-02  0.31481089 0.252 0.237 1.000000e+00
PPM1E  2.434700e-01 -0.04796266 0.110 0.119 1.000000e+00
```

```
FindMarkers(insulin_FGF2_AGG, ident.1 = "MG 2do I+F", ident.2 = "MG 3do I+F", features = c("DUSP1", "DUSP6", "PPP3CB", "PPP3R1", "PPM1A", "PPM1B", "PPM1D", "PPM1E", "PPM1G"), min.pct = 0.005, logfc.threshold = 0.02)
              p_val  avg_log2FC pct.1 pct.2    p_val_adj
DUSP1  3.697207e-26 -0.44818034 0.117 0.187 5.49368e-22
PPM1B  2.981483e-04 -0.15109036 0.080 0.100 1.00000e+00
DUSP6  3.129779e-03 -0.06534077 0.013 0.020 1.00000e+00
PPP3CB 2.772188e-02  0.03831759 0.030 0.023 1.00000e+00
PPM1E  4.356741e-02 -0.05383922 0.094 0.106 1.00000e+00
PPP3R1 1.041689e-01  0.02741809 0.065 0.057 1.00000e+00
```

```
FindMarkers(insulin_FGF2_AGG, ident.1 = "MG 2do I+F", ident.2 = "MGPCs", features = c("DUSP1", "DUSP6", "PPP3CB", "PPP3R1", "PPM1A", "PPM1B", "PPM1D", "PPM1E", "PPM1G"), min.pct = 0.005, logfc.threshold = 0.02)
              p_val  avg_log2FC pct.1 pct.2     p_val_adj
DUSP1  2.189300e-52 -0.55887356 0.117 0.237 3.253081e-48
PPM1B  1.208262e-11 -0.20488634 0.080 0.125 1.795357e-07
PPM1E  3.041839e-04 -0.09806454 0.094 0.119 1.000000e+00
PPM1D  3.860821e-03 -0.09088838 0.064 0.080 1.000000e+00
PPM1G  1.032669e-02 -0.07889363 0.086 0.102 1.000000e+00
DUSP6  3.923179e-02 -0.03568968 0.013 0.018 1.000000e+00
PPP3R1 3.972552e-02 -0.06253983 0.065 0.077 1.000000e+00
PPM1A  5.827332e-02 -0.05449119 0.034 0.042 1.000000e+00
```

**Statistics for dotplot figure 8f**

```
FindMarkers(clod_MG, ident.1 = "SS", ident.2 = "clod_SS", features = c("DUSP1", "DUSP6", "PPP3CB", "PPP3R1", "PPM1A", "PPM1B", "PPM1D", "PPM1E", "PPM1G"), group.by = "orig.ident", min.pct = 0.005, logfc.threshold = 0.05)
              p_val  avg_log2FC pct.1 pct.2 p_val_adj
DUSP6  0.0001184136 -0.25912643 0.143 0.208         1
PPP3R1 0.0070330894 -0.20446768 0.124 0.167         1
PPP3CB 0.0196159439 -0.19483543 0.200 0.241         1
DUSP1  0.0220592275 -0.19287668 0.167 0.204         1
PPM1A  0.1096064420  0.05441090 0.262 0.226         1
PPM1D  0.4162625077 -0.06098421 0.058 0.067         1
```

```
FindMarkers(clod_MG, ident.1 = "SS", ident.2 = "NM", features = c("DUSP1", "DUSP6", "PPP3CB", "PPP3R1", "PPM1A", "PPM1B", "PPM1D", "PPM1E", "PPM1G"), group.by = "orig.ident", min.pct = 0.005, logfc.threshold = 0.05)
              p_val  avg_log2FC pct.1 pct.2     p_val_adj
DUSP1  1.353880e-97 -1.42828753 0.167 0.588 2.852084e-93
DUSP6  2.877652e-21 -0.35630569 0.143 0.325 6.062062e-17
PPM1E  5.215232e-17 -0.29821011 0.119 0.271 1.098641e-12
PPP3R1 1.299999e-14 -0.31392090 0.124 0.267 2.738578e-10
PPM1G  6.138665e-07 -0.07273356 0.125 0.219 1.293171e-02
PPP3CB 1.273268e-06 -0.12940391 0.200 0.316 2.682267e-02
PPM1A  1.944819e-06  0.34549657 0.262 0.207 4.096957e-02
PPM1D  3.092502e-06 -0.10870423 0.058 0.117 6.514665e-02
```

```
FindMarkers(clod_MG, ident.1 = "NM", ident.2 = "clod_NM", features = c("DUSP1", "DUSP6", "PPP3CB", "PPP3R1", "PPM1A", "PPM1B", "PPM1D", "PPM1E", "PPM1G"), group.by = "orig.ident", min.pct = 0.005, logfc.threshold = 0.05)
              p_val  avg_log2FC pct.1 pct.2     p_val_adj
```

| | | | | | |
|---|---|---|---|---|---|
| DUSP1 | 5.276104e-40 | 1.16092182 | 0.588 | 0.270 | 1.111464e-35 |
| PPM1G | 2.108218e-05 | 0.22575692 | 0.219 | 0.132 | 4.441173e-01 |
| PPM1A | 1.859246e-03 | -0.21092452 | 0.207 | 0.260 | 1.000000e+00 |
| DUSP6 | 7.720548e-03 | 0.13350417 | 0.325 | 0.258 | 1.000000e+00 |
| PPP3R1 | 1.114018e-01 | 0.11970862 | 0.267 | 0.221 | 1.000000e+00 |
| PPM1B | 2.147841e-01 | -0.07253021 | 0.262 | 0.280 | 1.000000e+00 |
